# Generating ecological baselines on short-grass formations or “grazing lawns” in Central Terai landscape

**DOI:** 10.1101/2025.09.26.678694

**Authors:** Sankarshan Rastogi, Ashish Bista, Lovepreet Singh, Pranav Chanchani, T. Rengaraju, Naveen Khandelwal, Manish Singh, Mudit Gupta, H. Raja Mohan, Sanjay Pathak

## Abstract

Short-grass formations or the “grazing lawns” as they are popularly called support large congregations of mammalian herbivores globally. These systems are known to be maintained by cyclic process of grazing and nutrient deposition in the soil by foraging animals. The basic ecological characteristics of these short-grass systems have extensively been studied in the African context with little understanding particularly from the Himalayan foothills in Terai. In this study, we focused on understanding the ecological baselines such as grass species associations and associated nutrient content in these grazing lawns and tall grass formations across two protected areas of the Uttar Pradesh Terai in India. Our results show that, the protected areas had relatively similar grass species composition and associated nutrient content in the short-grass formations. However, when compared with tall grass formations, the grazing lawns differed markedly in both their grass associations and thus forage quality. Therefore, the results from this study indicate the need to maintain such grazing lawn habitats within the characteristic tall-wet grasslands of the Terai. This also indicates the importance of short-grass formations in providing palatable forage to the mammalian herbivores particularly in peak dry seasons of autumn and winters.

## Introduction

Grazing lawns are primarily short grass formations maintained primarily by positive feedback (Olff & Ritchie, 1998; Archibald, 2008; Hempson et al., 2015). These include intensive grazing by herbivores which keep the grass height at check typically <1 m and sometimes controlled fires (Karki et al., 2000; Donaldson et al., 2017). These short grasses provide highly nutritious forage favourable especially for small body-sized herbivores which are specialists in their diet selection (Wegge et al., 2006). In addition, the large congregations of mammalian ungulates also play an important role in maintaining these grassland habitats and regulating patterns of energy and nutrient flow in these system through grazing and defecation which enriches the soil (Milchunas et al., 1988; Hobbs, 1996; Olff, Ritchie & Prins, 2002). This has been true for grassland ecosystems globally which include African savannas, floodplain grasslands of Himalayan foothills and prairies of the North America (Dinerstein, 1980; Milchunas et al., 1988; Shaffer, 2019; Rastogi et al., 2022).

Our understanding about short-grass formations has majorly been through the extensive field-based research from the African savannas. These included studies on plant-herbivore interactions and their effects on such systems along with factors governing grazing in these grass formations by herbivores (McNaughton, 1984; McNaughton & Georgiadis, 1986; Milchunas et al., 1988). Though these studies laid the foundation in highlighting some relevant baselines about these important habitats, further research has majorly focused on emphasizing the nutritional parameters through experiment-based approach (Cromsigt & Olff, 2008; Zwerts et al., 2015). Nonetheless, not much is known about the short grass formations of the Himalayan foothills, the Terai which are otherwise limited in their spatial extent. These are majorly distributed in an interspersed manner within the tall grass formations which are densely clumped and often impenetrable (Ahrestani & Sankaran, 2016). This acts as a limiting factor in conducting proper field-based ecological surveys of these lawns which are otherwise an important habitat type for the herbivores especially the obligate ungulates (Owen-Smith et al., 2015). Our knowledge on these otherwise understudied grass formations has been derived from alluvial grasslands of Nepal Terai majorly focusing on their aspects of species diversity indices and nutrition parameters (Karki et al., 2000; Thapa et al., 2021). However, understanding the essential ecological facets such as species composition, forage quality, herbivory dynamics and environmental factors governing their occurrence among others is poorly known from both Indian and Nepal parts of the Terai Arc Landscape (TAL).

Hence, in this study we attempt to generate ecological baselines namely, vegetational characteristics within these short-grass formations. We attempt doing this at two levels, namely, i.) at the scale of protected area (Dudhwa Tiger Reserve & Pilibhit Tiger Reserve) collating data from all grassland plots together and ii.) at a small scale, where lawns differed by grass height within a particular protected area, i.e., lawns with a grass height > 1m and those with < 1m height respectively. To this end, we had specific objectives for these short-statured grasslands at the two levels of analysis; a) To understand the grass community structure and, b) Deciphering the forage quality of these lawns, across grassland habitats within DTR and PTR in the Indian Terai.

## Methods

### Study area

This study was conducted in the floodplain grasslands spanning two tiger reserves in the central Terai region of the Himalayan foothills in the state of Uttar Pradesh, India between January 2023-June 2023. These included grassland habitats within Pilibhit Tiger Reserve (PTR) and in and around two protected areas namely, Dudhwa National Park & Kishanpur Wildlife Sanctuary which otherwise, together with Katarniaghat Wildlife Sanctuary form the Dudhwa Tiger Reserve (DTR). Both tiger reserves are separated by river Sharda, a tributary of Ganga, which originates from upper reaches of Uttarakhand straddling along western Nepal south of which lies its floodplains while entering the district of Pilibhit and eventually Lakhimpur Kheri in Uttar Pradesh. The hydrodynamics of Sharda and its discharge in the form of reservoir and linear canal systems plays a crucial role in rejuvenating the interspersion of moist-deciduous Sal forests, riverine forests, grasslands, and wetland habitats of this conservation ecoregion (Olson & Dinerstein, 2002; Mathur & Midha, 2008). Collectively, DTR and PTR support one of the largest expanses of the floodplain grassy biomes in the TAL. These are mostly dominated by the tall-clumped grass formations of *Themeda arundinacea, Narenga porphyrocoma, Saccharum spontaneum, Saccharum bengalense* and *Sclerostachya fusca* embedded within which are patches of nutrient-rich short grasses such as *Imperata cylindrica, Apluda mutica, Cynodon dactylon, Dicanthium annulatum* and *Vetiveria zizanioides*.

These habitats are vital for several threatened grassland obligate species birds and mammalian herbivores including swamp francolin (*Francolinus gularis*), Bengal florican (*Houbaropsis bengalensis*), hispid hare (*Caprolagus hispidus*), one-horned rhinoceros (*Rhinoceros unicornis*), Asiatic elephant (*Elephas maximus*), hog deer (*Axis porcinus*) and swamp deer (*Recurvus duvaucelii duvaucelii*). However, the declining trends in populations of these species is majorly attributable to the substantial loss of grasslands due to conversion into agricultural lands owing to the highly productive edaphic conditions, habitations to support dense human populations and other infrastructural purposes (Strahorn, 2009; Chanchani et al., 2014).

### Study design

In order to generate information on vegetation attributes of the short-grass formations or grazing lawns, as a first step, we conducted reconnaissance surveys to delineate and spatially map such formations in and around the study area. The relevant structuring of spatial grassland data sourced from Chanchani, 2016, ground-truthing verifications and pilot surveys in the study area were done as part of the larger long-term grassland ecology, monitoring and conservation project initiated by WWF-India in collaboration with Uttar Pradesh Forest Department between March, 2021-February, 2022. Given, ecological succession of these grasslands into Sal climax community is an imminent phenomenon in the Terai (Kumar et al., 2002; Shukla, 2009; Chanchani et al., 2014), the boundaries for some of the short-grass formations were readjusted and some additional lawns digitized using Quantum GIS (QGIS) version 3.16.2 (QGIS Development Team, 2022) based on ground-truthing data.

As a next step, the short-grass formations were broadly categorized based on their grass species height into those with grass stature > 1m and those with < 1m height. We characterized vegetation attributes of the identified lawns in a sampling unit of one-hectare plots (1-ha hereafter). These were selected following the reasons stated in Rastogi et al., 2022. We sampled nearly equal number of plots in the grassland formations of the two tiger reserves differentiated by grass height (Table 1).

**Table 1:** Classification of sampling plots across two tigers reserves and grass height.

| **Tiger Reserve** | **Grass height type** | **Number of 1-ha plots** |
| --- | --- | --- |
| Dudhwa (n=38) | > 1m | 17 |
|  | < 1m | 21 |
| Pilibhit (n=37) | > 1m | 21 |
|  | < 1m | 16 |

### Field data collection protocols

We employed slightly varied yet similar field data collection methods in the two PAs.

In Dudhwa Tiger Reserve, we collected data on vegetation attributes of grasses along four equally spaced (20 m) transects, T1-T4 within each 1-ha sampling plot using point-intercept method as depicted in figure 2 (Evans & Love, 1952; McNaughton, 1983; Rastogi et al., 2022). At a 10 m interval along each of the transect, we noted down the grass species that hit the tip of surveyors’ shoes. Altogether, 40 intercepts were sampled within each 1-ha plot (10 points per transect). The same survey design was used for the grass formations across grass height types. Moreover, in order to understand the forage quality of the short-grass formations, we sourced C:N ratio values for each of the dominant grass species from Rastogi et al., 2022.

**Figure 1:**
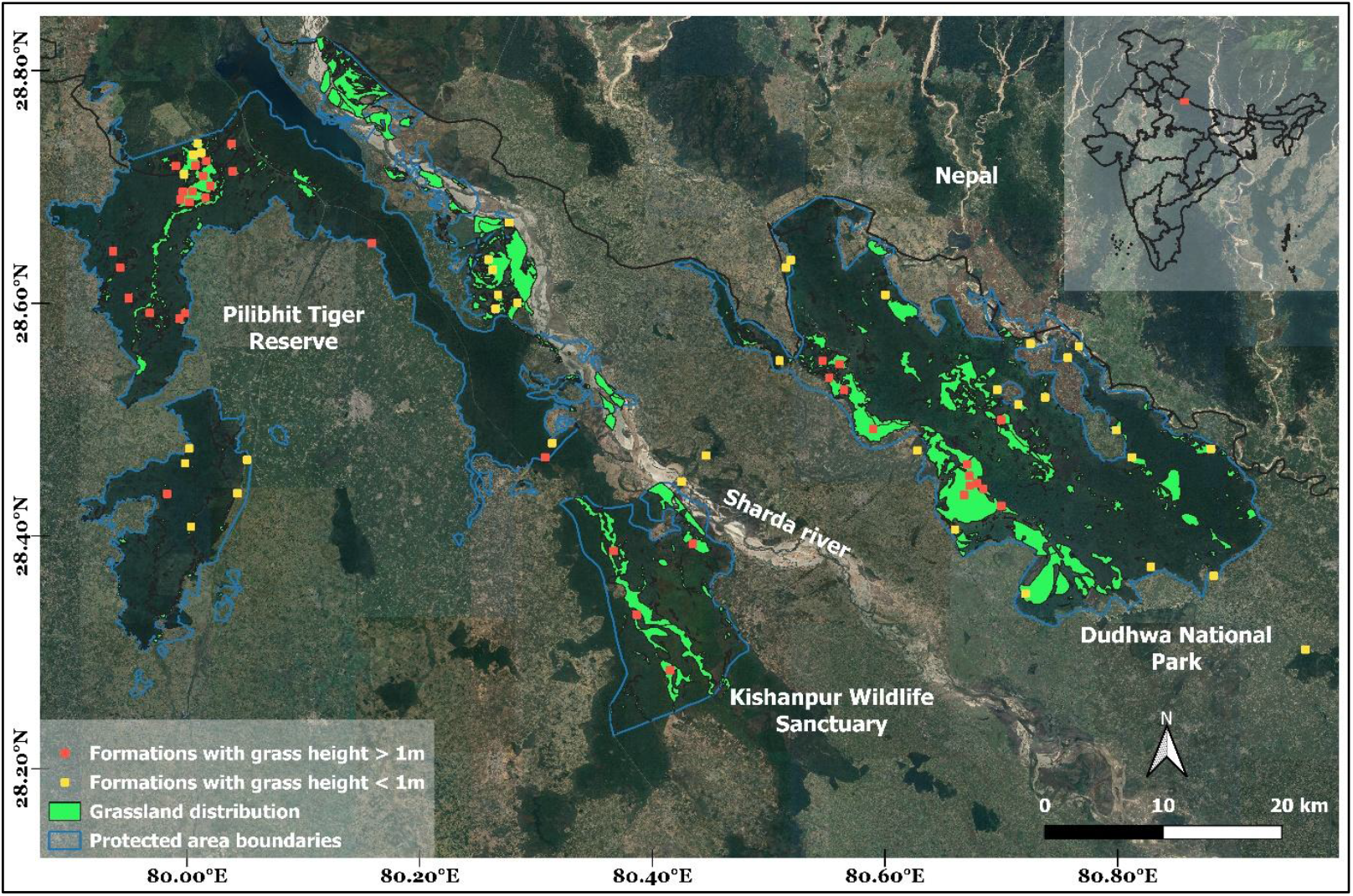
Spatial distribution of sampling plots across two tiger reserves of Central Terai in India.

**Figure 2:**
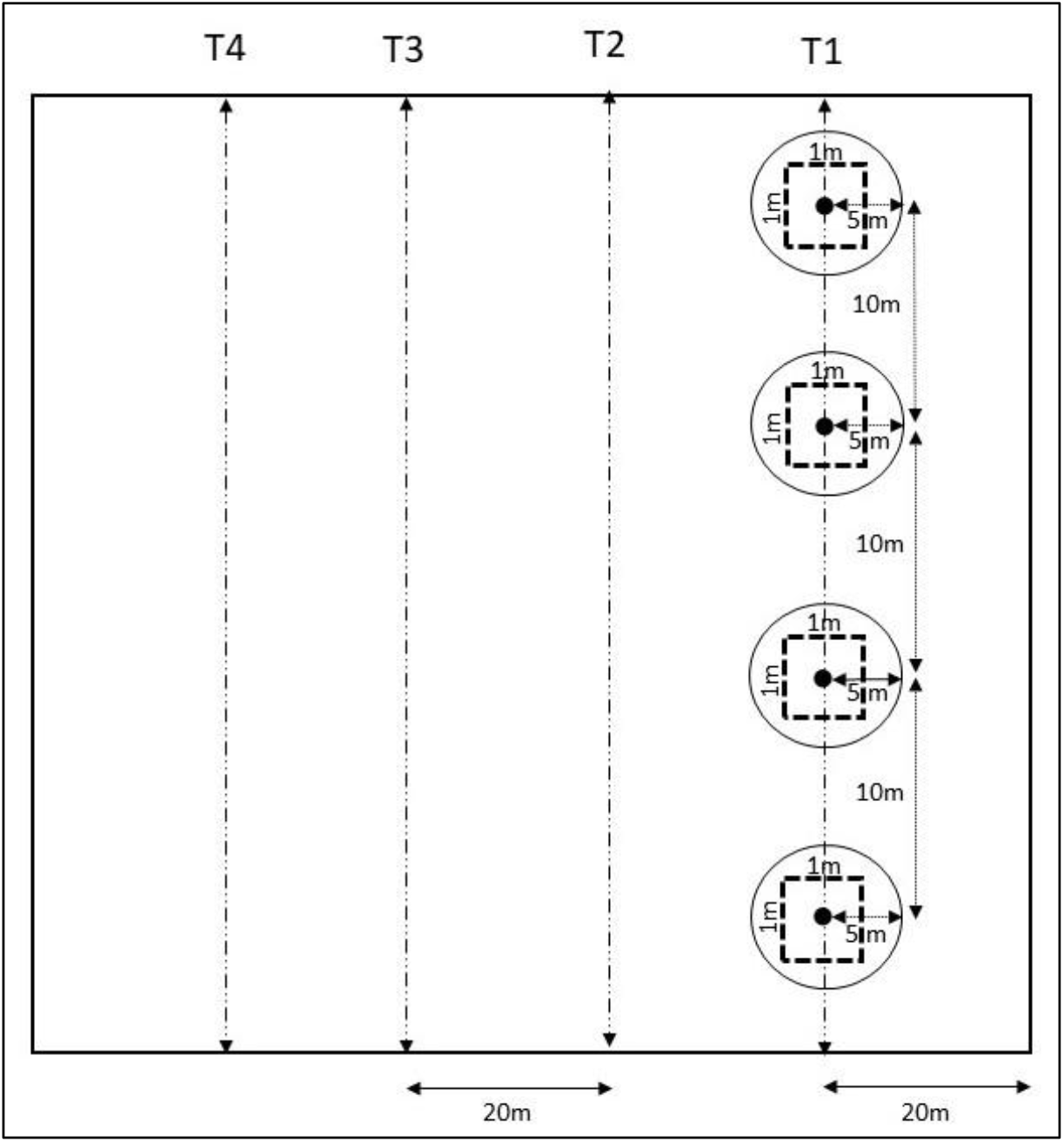
Schematic of sampling plot with transects and representative sampling intercepts (10 m replicate) along which data on vegetation characteristics was collected.

However, in Pilibhit Tiger Reserve, we followed the same field data collection protocol as used in Rastogi et al., 2022 since there was a need to generate relevant baselines for grasslands in this area as per the instructions from respective management authorities. Under this framework, within the 1-ha sampling plot, we walked three transects separated by 20 m and each point-intercept was separated by 5 m along which the data on grass species ID was collected (Rastogi et al., 2022).

### Data Analysis

We analyzed the data on vegetation attributes, namely grass community composition and forage quality at two spatial scales –

1. At the level of individual Protected area
2. 1-ha plots differentiated by grass height in each Protected area

### Grass community analysis

Cluster analysis was used to identify grass communities in the short-grass formations across two grazing types for each of the protected area as mentioned in table 1. We used hierarchical agglomerative clustering using the function *hclust* in *vegan* package of R to classify the sampling plots into discrete groups based on the relative abundance of various grass species in the 1-ha plot (Oksanen et al., 2013). The ecological distance between the clusters was determined using Bray-Curtis dissimilarity method of vegetation composition (Beals, 1984). We used “complete” as the clustering method in order to generate different clusters since it uses pairwise dissimilarities between different elements (1-ha plots here), and considers the largest value of the dissimilarities as the distance between two clusters. The resulting clusters (grass communities here) were determined using the threshold as the number of nodes emerging out from the primary branch of division in the final cluster. In order to assign an identity to each community, relative abundance of each grass species was computed as the number of hits (intercepts) for a particular species/the total number of hits for all species in the respective cluster/community. Further, the species with highest abundance were used as the representative of the individual grass community.

### Forage quality analysis

We used C:N ratios (proxy for nutrition values) for the dominant grass species as determined using laboratory-based analysis reported in Rastogi et al., 2022. As a first step in this analysis, for each sampling plot, we computed the community weighted C:N ratio by multiplying C:N ratio for a particular grass species with their relative abundance in a respective 1-ha plot. For the grass species whose C:N ratio data was not reported in Rastogi et al., 2022, we assigned them the community weighted mean value of the 1-ha plot as generated using ∼90% of the grass individuals that formed the respective plot (Osuri et al., 2020; Rastogi et al., 2022). C:N ratios for each grass community type across the two grazing categories were calculated by averaging the weighted C:N ratios for all plots associated to that community, and given low number of sampling plots in some communities, standard errors were estimated by bootstrapping (Efron & Tibshirani, 1985; Osuri et al., 2020).

## Results

Overall, across the two PAs, we recorded 19 grass species in the short-grass formations with differences in the species type. Three grass species namely, *Imperata cylindrica, Vetiveria zizanioides, Cynodon dactylon* accounted for about 41% and 46% of the total grass abundance across the sampling plots in the two PAs respectively. In addition, *Narenga porphyrocoma* (12%) in DTR and *Desmostachya bipinnata* (11%) in PTR contributed to the total grass abundance in the study area.

### At the scale of each Protected area

The short-grass formations in Dudhwa Tiger Reserve were dominated by three grass communities: *Imperata cylindrica* – *Vetiveria zizanioides* (IC-VZ), *Narenga porphyrocoma* – *Imperata cylindrica* (NP-IC) and *Cyperus kyllingia* - *Cynodon dactylon* (CK-CD) (Figure 3). Community weighted C:N ratios, used as an inverse surrogate for forage quality differed for the three communities where *Cynodon-Cyperus* community had the highest palatability (since C:N ratios were lowest) followed by the two other communities which in comparison had a lower palatability for the herbivores (Figure 3).

**Figure 3:**
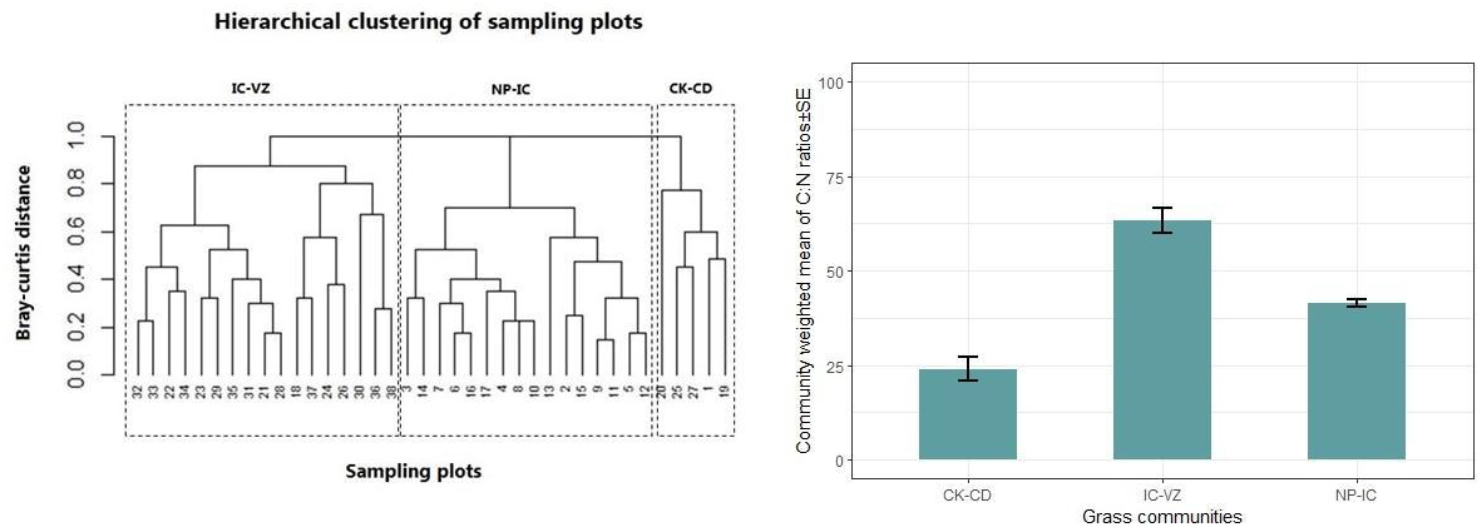
Dendrogram depicting the overall grass community structure and community weighted C:N ratios for the three identified communities in Dudhwa Tiger Reserve.

Similarly, grazing lawns in Pilibhit Tiger Reserve clustered into three grass communities: *Imperata cylindrica* – *Desmostachya bipinnata* (IC-DB), *Cynodon dactylon* – *Imperata cylindrica* (CD-IC) and *Imperata cylindrica* – *Vetiveria zizanioides* (IC-VZ) (Figure 4). In this case, *the Cynodon-Imperata* community had the highest palatability (lowest C:N ratio value) while two *Imperata* communities were marginally less palatable (Figure 4).

**Figure 4:**
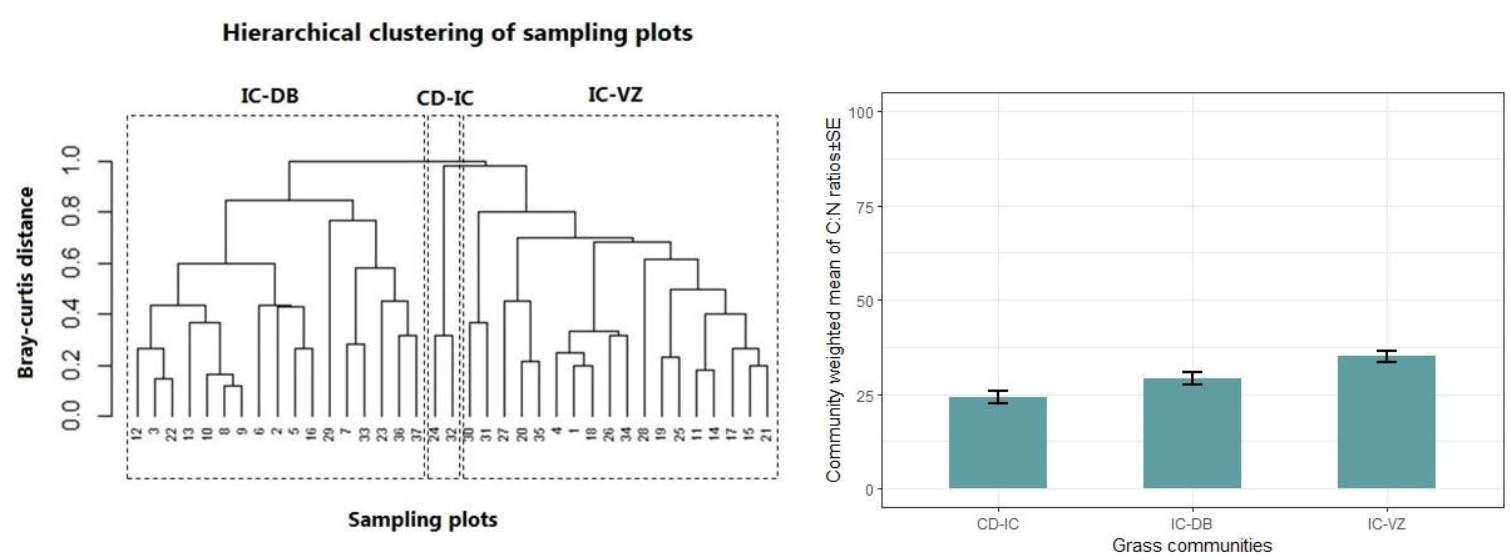
Dendrogram depicting the overall grass community structure and community weighted C:N ratios for the three identified communities in Pilibhit Tiger Reserve.

### At the grass height scale within each Protected area

#### 1. Dudhwa Tiger Reserve

When the 1-ha plots were categorized by grass height into a) plots with grass height > 1m and b) plots with grass height < 1m, there were certain differences in the grass communities dominating each grazing type. The plots predominantly with grass height > 1m had three grass communities: *Narenga porphyrocoma* – *Imperata cylindrica* (NP-IC), *Imperata cylindrica* – *Desmostachya bipinnata* – *Narenga porphyrocoma* (IC-DB-NP) and *Sclerosatchya fusca* – *Cyperus kyllingia* (SF-CK) (Figure 5). Interestingly, these three communities had high community weighted C:N ratios indicating overall low palatability in such short-grass formations.

**Figure 5:**
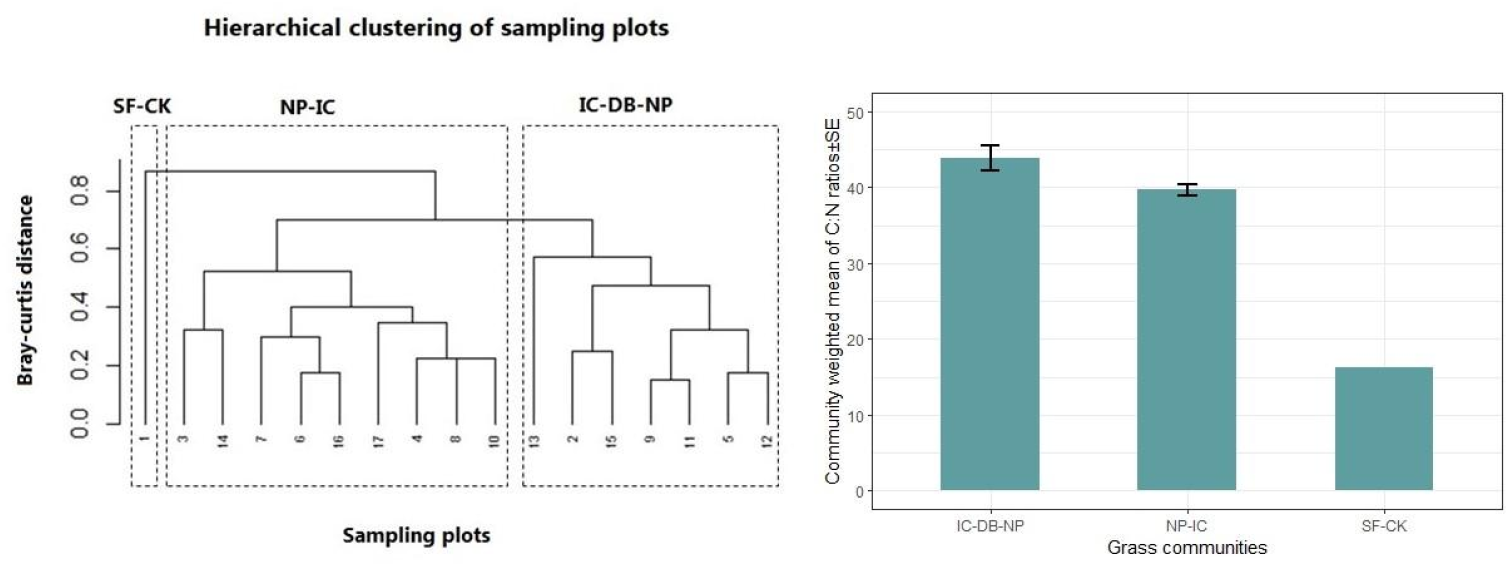
Dendrogram depicting the grass community structure and community weighted C:N ratios for the three identified communities within the > 1m grass height plots in Dudhwa Tiger Reserve.

On the other hand, the short-grass formations with grass height < 1m had different grass communities: *Cyperus kyllingia* - *Cynodon dactylon* (CK-CD), *Imperata cylindrica* – *Cynodon dactylon* (IC-CD) and *Hemarthria compressa* – *Vetiveria zizanioides* (HC-VZ) (Figure 6). Among these, *the Cyperus-Cynodon* community had the highest nutritive value than the other two communities (Figure 6).

**Figure 6:**
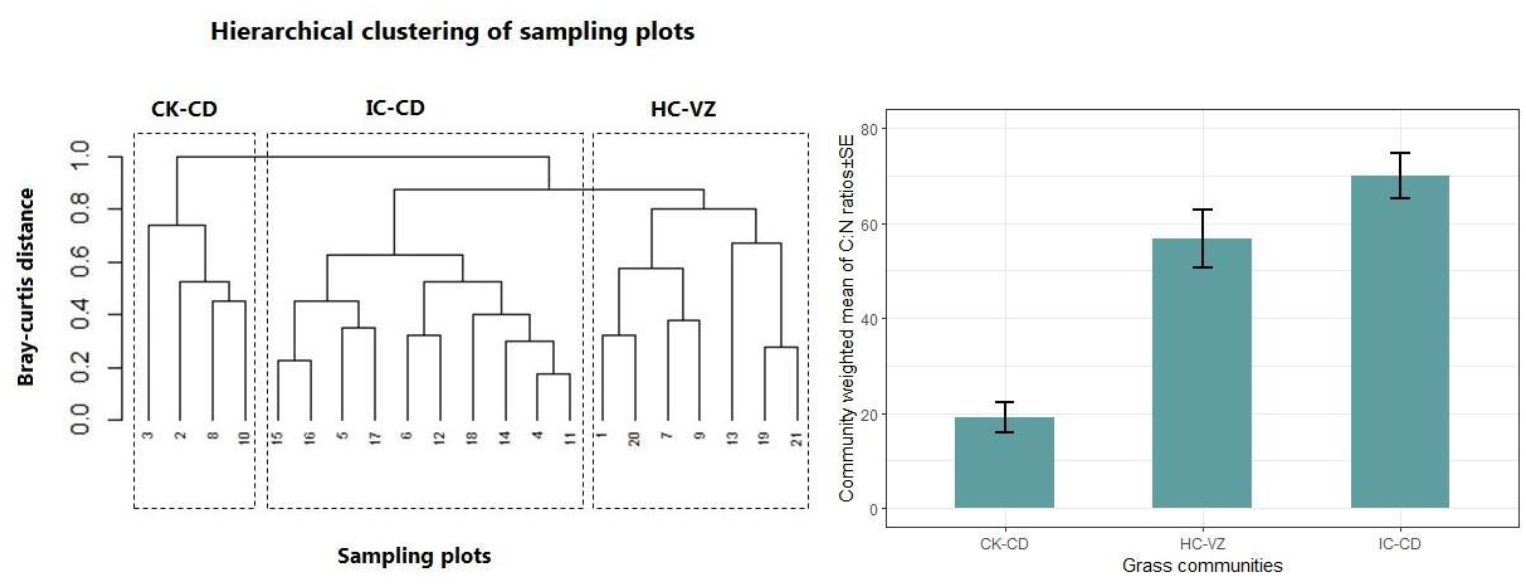
Dendrogram depicting the grass community structure and community weighted C:N ratios for the three identified communities within the < 1m grass height plots in Dudhwa Tiger Reserve.

#### 2. Pilibhit Tiger Reserve

The short-grass formations in this protected area revealed three grass communities each for the two grass height types with differences in species associations within the community. The 1-ha plots with > 1m grass height had: *Cynodon dactylon* - *Vetiveria zizanioides* (CD-VZ), *Imperata cylindrica* – *Cymbopogon jwarancusa* (IC-CJ) and *Desmostachya bipinnata* – *Imperata cylindrica* (DB-IC) as the dominant grass communities. However, the former grass community was only present in one of the sampling plots (thus, missing standard error bars) and had a higher palatability than the latter two community associations with *Imperata cylindrica* (Figure 7).

**Figure 7:**
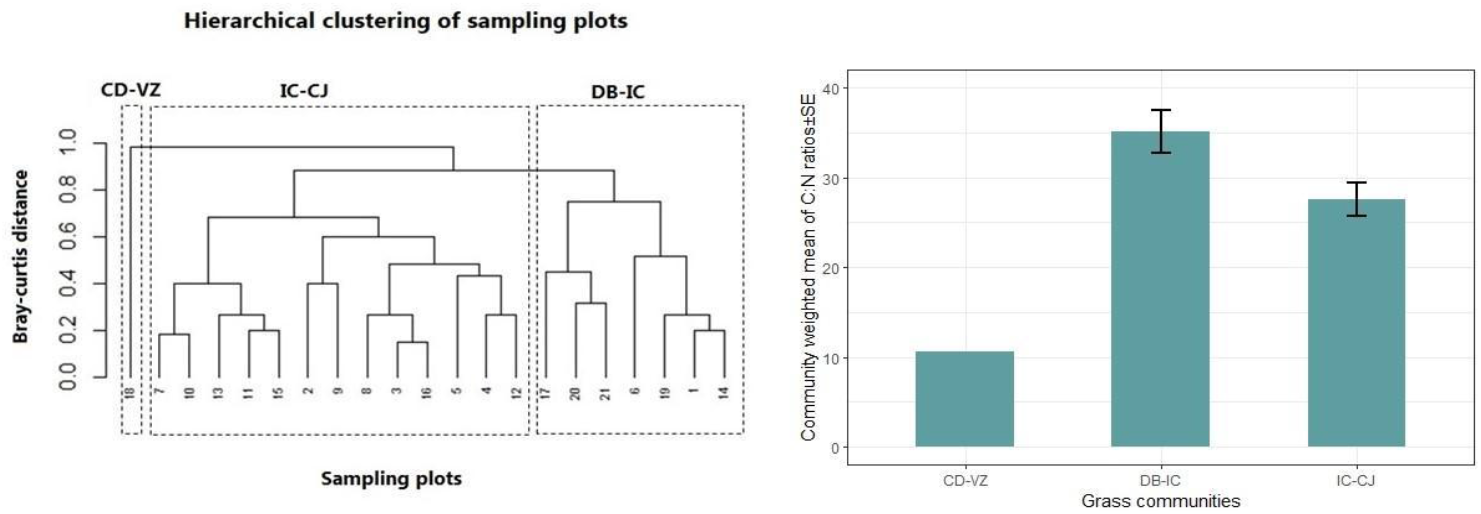
Dendrogram depicting the grass community structure and community weighted C:N ratios for the three identified communities within the > 1m grass height plots in Pilibhit Tiger Reserve.

In the case of plots < 1m grass height, *Cynodon dactylon* - *Vetiveria zizanioides* (CD-VZ), *Imperata cylindrica* – *Saccharum spontaneum* (IC-SS) and *Imperata cylindrica* – *Cynodon dactylon* (IC-CD) formed the dominant grass communities in this grass height type. Forage quality was higher for the *Cynodon-Vetiveria community* than the other two *Imperata* dominant associations (Figure 8).

**Figure 8:**
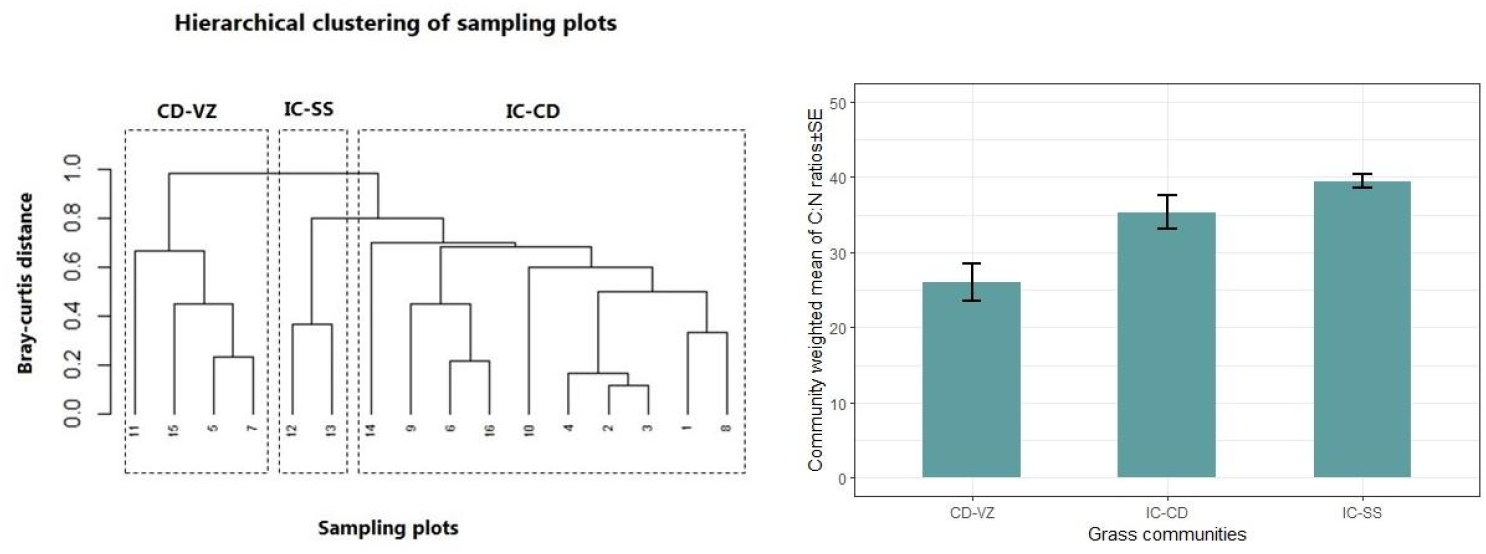
Dendrogram depicting the grass community structure and community weighted C:N ratios for the three identified communities within the < 1m grass height plots in Pilibhit Tiger Reserve.

## Discussion

The short grass formations of the Terai act as important grazing habitats for the mammalian herbivores in terms of their dietary needs (Karki et al., 2000). These habitats are crucial for small and medium-sized herbivores such as hog deer, wild boars and swamp deer which are foregut fermenters and tend to depend on quality of forage than quantity (Wegge et al., 2006; Ahrestani et al., 2009). Therefore, understanding the ecological characteristics of small grass formations is essential to sustain the populations of such endemic herbivores. Moreover, they are also important for monogastric and other large megaherbivores like elephant, rhino and wild buffalo during season of scarcity.

The results from this study across two important tiger reserves of Central Terai indicate nearly similar grass community structure across two grass height types, i.e. > 1m and < 1m plots. Consistent with the findings of the previous research in the Terai landscape, *Cynodon dactylon* dominated grass communities tend to be richer in forage nutrition than others across the grazing types (Thapa et al., 2021; Rastogi et al., 2022). The study also highlights the contribution and importance of *Cyperus* spp. in the enrichment of grasslands with their high palatability. Family Cyperaceae, which is an integral part of grassland ecosystem, had been earlier ignored. These results are also important in the context of the necessary management interventions essential to sustain the overall grassland habitats of the Gangetic floodplains distributed across India and Nepal. Moreover, this data also establishes the information baseline on the status of grassland and will play indispensable role in long term monitoring of various ecological attributes.

Although, tall grasslands have their own significance in providing hide to tigers, forming sole habitat matrix for endangered mammals like hispid hare and food to innumerable granivore birds, but in the context of herbivores are poor in forage quality especially in the dry months between November – February. During this time, the height of grasses is at their maximum with low nutrient content (Olff et al., 2002; Rastogi et al., 2022). It must be noted that the equilibrium between tall and short grasses effectively earmarked by megaherbivores in the past has been now disturbed due to extermination of species like elephant, rhino and buffaloes from a major extent of these habitats. Thus, it is necessary to ensure year-round nutrient rich forage for specialist herbivores such as swamp deer and hog deer by maintaining a balance with high proportion of grazing lawns. Therefore, generating these ecological baselines will aid in better management of such habitats by implementing necessary interventions in a timely manner which will in turn ensure forage for the herbivores for a longer duration in the year in these otherwise threatened habitats of Terai.

Therefore, future work should focus on designing research studies which should incorporate other ecological factors such as edaphic characteristics, successional trends in grass vegetation, association and dynamics of non-poaceae herbaceous plants, herbivore distribution and influence of hydrological features in such short grass formations and how they differ from the typical tall grass formations of the Terai. Moreover, understanding the retrogression in community through experimental allowing of cattle grazing in undisturbed grasslands and vice-versa in controlled areas is the need of the hour. In future direct observational studies on herbivore feeding behaviour to substantiate laboratory analysis based upon C:N ratios to determine palatability should be marked. Absolute grade of pasturage quality of a grass also depends upon its potential of biomass production in a given time period. Hence, palatability and composition of a grassland must also be studied in association of total biomass yield potential.

## Acknowledgements

This study was part of the ongoing long-term grassland research, conservation and management in Dudhwa Tiger Reserve. We would like to thank the PCCF(Wildlife)/CWLW, Uttar Pradesh for granting research permits. We extend our sincere thanks to all the frontline forest staff of Dudhwa Tiger Reserve and Pilibhit Tiger Reserve for help in the field data collection for this study. At WWF-India, we would like to thank Mr. Ravi Singh, Dr. Sejal Worah, Dr. Dipankar Ghose, Mr. Yash Shethia, Mr. Saurav De, and Dr. Anil Kumar Singh for their constant guidance and support throughout the project. We also acknowledge the support of WWF-India’s TAL-UP team and our field collaborators, Mr. Ram Lakhan and Mr. Ram Bharose ji for their exceptional field knowledge and contributing in data collection.

